# The lysosomal glutamine transporter SLC38A7/SNAT7 modulates SAMHD1 antiviral activity and promotes HIV-1 production in human macrophages

**DOI:** 10.64898/2026.03.06.709337

**Authors:** F Herit, G Lê-Bury, Q Provôt, D To-Puzenat, J Haagen, Matozo de Souza T, A Dumas, M Morel, F Margottin-Goguet, C Sagné, A Sáez-Cirión, F Niedergang

## Abstract

HIV-1 (Human Immunodeficiency Virus type 1) infects macrophages, which resist to the cytopathic effects of the virus and are considered as viral reservoirs. However, the cellular factors involved in viral production by human macrophages have not been fully identified. In this study, we focused on the amino acid transporter SNAT7 (small neutral amino-acid transporter 7), member of the SLC38 solute carrier family, which is the main lysosomal transporter of glutamine from the lysosome to the cytoplasm. Its expression was increased by HIV-1 infection. We revealed that the absence of SNAT7 inhibited viral production not only at the level of protein synthesis, but also early at the level of reverse transcription, without affecting global RNA or protein synthesis in the cells. The reduction in HIV expression upon SNAT7 depletion correlated with a reduction in the levels of an inactive form of the SAMHD1 (SAM domain- and HD domain-containing protein) restriction factor and was rescued following SAMHD1 degradation. Lastly, supplementation of the extracellular medium with glutamine in the absence of SNAT7 partially restored viral production.

Together, our data reveal that glutamine extracted from lysosomes is involved in the early stages of the HIV-1 cycle and that the SNAT7 glutamine transporter acts as a dependency factor for HIV-1 in human macrophages.

## Introduction

Human immunodeficiency virus type 1 (HIV-1) is a lentivirus that infects immune cells and leads to acquired immunodeficiency syndrome (AIDS) characterized by the loss of CD4+ T cells, high levels of inflammation, and the development of opportunistic diseases. HIV-1 enters into its target cells through the CD4 receptor and CXCR4 or CCR5 co-receptor, which are expressed at the surface of CD4+ T cells, dendritic cells and macrophages (Deeks et al., 2015). We have previously shown that HIV-1 infection of human monocyte-derived macrophages (hMDM) disrupts key functions of these cells, such as phagocytosis, degradative properties, and cytokine secretion, facilitating the development of opportunistic bacteria (Dumas et al., 2015; Le-Bury and Niedergang, 2018; Mazzolini et al., 2010).

In contrast to CD4+ T cells, macrophages are resistant to the cytopathic effects of the virus and survive for long periods of time, continuously producing the virus (Koppensteiner et al., 2012). Indeed, together with CD4+ memory T cells, macrophages represent a viral reservoir hidden in tissues with restricted access to anti-retroviral treatment (ART) (Arainga et al., 2017; Castellano et al., 2017; Ganor et al., 2019; Kruize and Kootstra, 2019; Wong et al., 2019).

In lymphocytes, virions bud and are released from the plasma membrane (Ono, 2009), whereas in macrophages, HIV-1 assembly and budding occur at the membrane of a non-acidic compartment called the virus-containing compartment (VCC). This compartment is transiently connected to the plasma membrane, allowing for virus release in the extracellular medium (Benaroch et al., 2010; Gaudin et al., 2013; Jouve et al., 2007; Rodrigues et al., 2017; Tan and Sattentau, 2013).

However, the cellular factors involved in virus production by human macrophages have not yet been fully identified. Various steps of the viral cycle can be inhibited by host cell restriction factors, including SAM domain- and HD domain-containing protein (SAMHD1) that restrains HIV-1 replication in macrophages (Bowen et al., 2022). In addition, several studies previously showed that the metabolism of macrophages is modulated by HIV-1 (Castellano et al., 2019; Datta et al., 2016; Huang et al., 2011; Real et al., 2022; Saez-Cirion and Sereti, 2021). In this context, the solute carrier proteins (SLC) superfamily that comprises at least 450 proteins located on the cell or organelles’ surface, delivering a wide array of solutes, including amino-acids, through membranes, is of particular interest. SLC proteins are involved in the metabolism of immune cells (Cesar-Razquin et al., 2015; Pizzagalli et al., 2020; Song et al., 2020). SLC were shown to be modulated in immune cells upon HIV-1 infection. For instance, the alanine transporter SNAT1 is down-regulated by HIV-1 Vpu to reduce T cells proliferation (Matheson et al., 2015). By contrast, other SLC were described to be dispensable for cell survival, but necessary for virus expansion, constituting HIV dependency factors (Ali et al., 2021; Montoya, 2023; Park et al., 2016).

As macrophages are professional phagocytes with highly degradative properties (Depierre et al., 2023; Fairn and Grinstein, 2012), we considered the potential involvement of endo-lysosomal transporters in HIV-1 production. We focused on the major lysosomal transporter of glutamine called SLC38A7/SNAT7, which exports glutamine from the lysosomal lumen to the cytoplasm (Verdon et al., 2017). Glutamine is the most abundant amino acid in the blood and is actively involved in many biosynthetic and regulatory processes notably in leucocytes (Curi et al., 2005; Yoo et al., 2020, Cruzat et al., 2018; Newsholme, 1999). Moreover, glutamine metabolism is altered during HIV infection and supports viral replication (Clerc et al., 2019; Hegedus et al., 2017).

In this study, we showed that SNAT7 is preferentially expressed in macrophages, compared to monocytes and lymphocytes. Its protein expression is increased upon HIV-1 infection. In contrast, depletion of SNAT7 resulted in a decrease in HIV-1 production already at the level of reverse transcription. In addition, the antiviral activity of the HIV-1 restriction factor SAMHD1 could play a role along SNAT7. Our results reveal that SNAT7 and the glutamine from lysosomes are crucial for the early steps of HIV-1 production.

## Results

### SNAT7 is enriched in macrophages and increased upon HIV-1 infection

SNAT7 has been previously described to be expressed in mouse neurons and various cancer cells (Audet-Delage et al., 2023; Hagglund et al., 2011; Haratake et al., 2021; Meng et al., 2022; Verdon et al., 2017), but its expression in immune cells was unclear. Therefore, we set out to compare SNAT7 protein expression in macrophages and other immune cells targeted by the virus. Monocytes were isolated from fresh human blood and differentiated into macrophages, while T lymphocytes isolated in parallel were stimulated with cytokines. As shown in Fig. 1A, SNAT7 was induced after differentiation of monocytes into macrophages (Fig. 1B). In contrast, it was barely detectable in circulating or activated T lymphocytes. Similar differences were observed for SNAT7 RNA levels in the different cell types (Fig. 1C).

**Figure 1.**
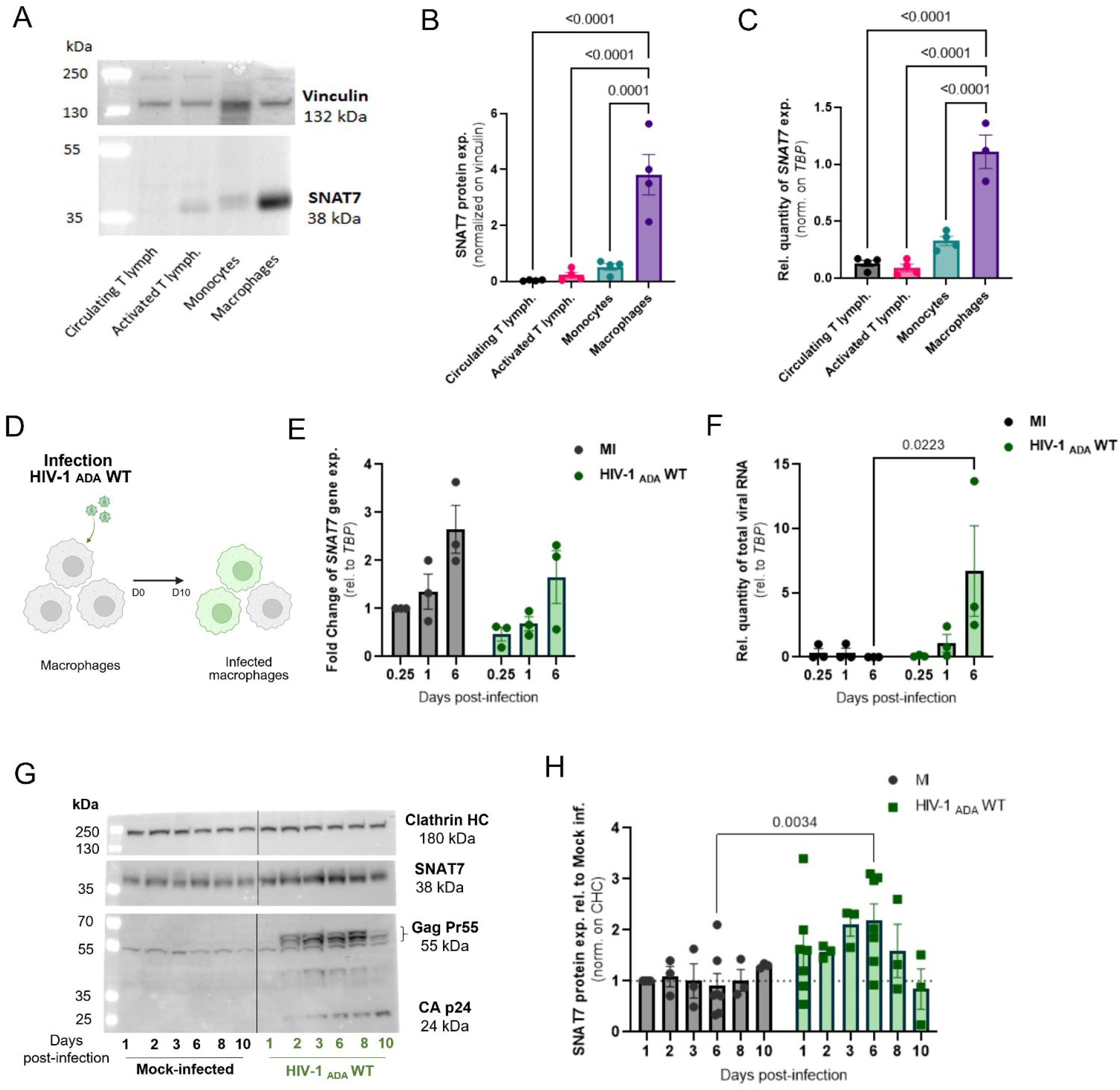
The glutamine transporter SLC38A7/SNAT7 is predominantly expressed by human macrophages, and its protein expression is increased upon HIV-1 infection. (A-C) Monocytes and lymphocytes were negatively isolated from peripheral blood mononuclear cells (PBMCs) of different donors using STEMCELL™ Technologies enrichment kits. Monocytes were differentiated into macrophages, and lymphocytes were activated with IL2 and PHA-P, for 6 d. Cells were lysed to collect proteins and RNA. (A) Representative immunoblot of SNAT7 protein expression by the different cell types, in parallel with vinculin as loading control. (B) Quantification of immunoblots of SNAT7 protein expression normalized on vinculin expression. (C) Relative quantity of *SNAT7* gene expression normalized on *Tata box protein* (*TBP*) calculated using the 2 ^(-ΔCt)^ method. Points represent the mean of 3 to 4 independent experiments +/- SEM. Ordinary one-way ANOVA statistical analysis was performed, the numbers above the graphs represent the p-value. (D-H) HMDM were infected with the HIV-1_ADA_WT strain or supernatant without particles (mock-infected, MI) from 1 to 10 d, and cells were lysed to collect RNA or proteins. (D) Experimental protocol. (E-F) HMDM were infected with the HIV-1_ADA_WT strain or mock-infected for 0.25 (i.e. 6 h) to 6 d, and cells were lysed to collect RNA. (E) Fold change in *SNAT7* gene expression normalized on *TBP* calculated using the 2 ^(-ΔΔCt)^ method. (F) Relative quantity of HIV-1 gene expression normalized on *TBP* calculated using the 2 ^(-ΔCt)^ method. Points represent the mean of 3 independent experiments +/− SEM. Two-way ANOVA statistical analysis was performed, the numbers above the graphs represent the p-value when p < 0.05. (G) Representative immunoblot of SNAT7, HIV-1 Gag Pr55 and CAp24 protein expression at the indicated time points, in parallel with Clathrin HC as loading control. (H) Quantification of immunoblots of SNAT7 protein expression normalized on Clathrin HC. Points represent the mean of 3 independent experiments +/− SEM. Two-way ANOVA statistical analysis was performed, the numbers above the graphs represent the p-value when p < 0.05.

To analyze if SNAT7 expression is modulated upon HIV-1 infection, human macrophages were infected with the HIV-1_ADA_WT strain (Fig. 1D). *SNAT7* RNA level was quantified from 0.25 (i.e., 6 h) to 6 days post-infection, revealing an increase of gene expression during the time course of cell differentiation both in mock- and in HIV-1-infected cells (Fig. 1E and Supp. Fig 1A). HIV-1 viral genes were detected since day 1 post-infection and became abundant after 6 days compared to mock-infected cells (Fig. 1F and Supp. Fig. 1B). In contrast, SNAT7 protein expression was increased from 1 day to 8 days post-HIV-1 infection compared to control cells, before returning to basal levels 10 days post-infection (Fig. 1G, H).

In conclusion, SNAT7 protein expression is a macrophage-specific signature that is rapidly and transiently increased and/or stabilized during HIV-1 infection.

### SNAT7 is necessary for an efficient production of HIV-1 by human macrophages

To evaluate a potential role of SNAT7 on HIV-1 infection, hMDM were infected or not (Mock) with HIV-1_ADA_WT and treated with 2 different sequences of siRNA targeting SNAT7 mRNA (#1 and #2), or with control (Ctl) siRNA (Fig 2A, B). SNAT7 protein expression, as determined by immunoblotting, was decreased by siRNA treatment from 30 to 60 % compared to siRNA-treated control cells (Fig 2C). SNAT7 depletion induced a reduction of GagPr55 expression by 55 % and 73 % for SNAT7 siRNA #1 and #2, respectively, compared to cells infected in control conditions (Fig. 2D). Strong reductions, above 85 %, were also observed in Cap24 production, both in cell lysates and in supernatants (Fig 2E, F), as well as in the release of infectious viral particles (Fig. 2G). Of note, similar results were obtained using VSVG-pseudotyped HIV-1_ADA_WT virus (Supp. Fig 2A-D), showing that the role of SNAT7 in viral production does not depend on the way the virus enters in macrophages during the initial infection. Importantly, the impact of siRNA treatment on viral infection was not related to cytotoxicity, measured by lactate dehydrogenase (LDH) release in the supernatant, (Fig. 2H), nor to changes in the cellular concentrations of RNA and protein extracted from each condition (Supp. Fig. 2E, F).

**Figure 2.**
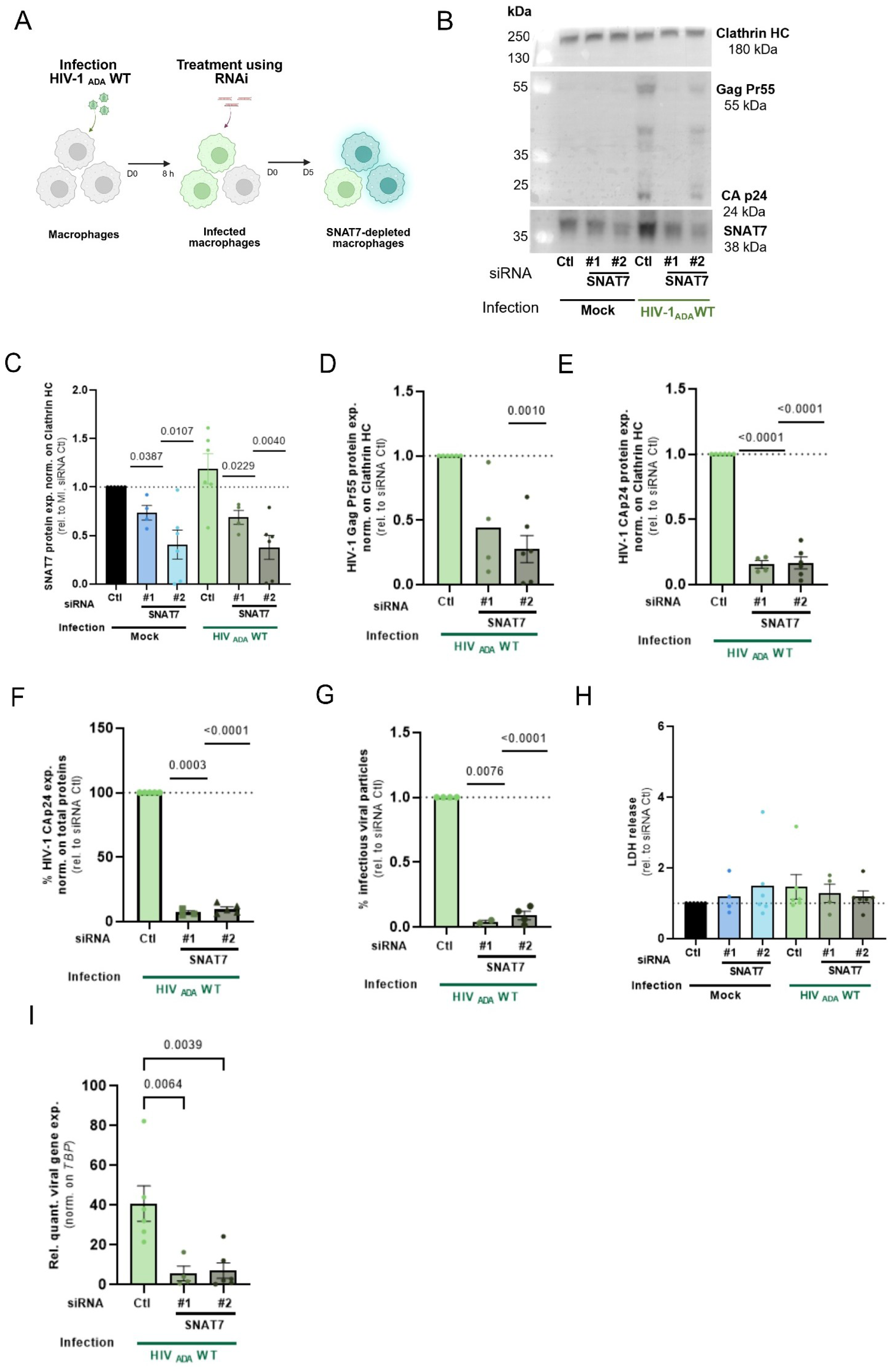
SNAT7 is necessary for the efficient production of HIV-1 by human macrophages. (A-I) Human macrophages were infected with the HIV-1_ADA_WT strain or mock-infected for 8 h, washed and treated with siRNA duplexes targeting SLC38A7/SNAT7 or luciferase encoding gene as control. Macrophages were cultured for 5 d, supernatants were stored and cells were lysed to collect proteins. (A) Experimental scheme to obtain samples analyzed in (B-I). (B) Representative immunoblot of SNAT7, HIV-1 Gag Pr55 and CAp24 protein expression, in parallel with Clathrin HC as a loading control. (C) Quantification of immunoblots of SNAT7 protein expression normalized to Clathrin HC. (D) Quantification of immunoblots of HIV-1 Gag Pr55 protein expression normalized to Clathrin HC. (E) Quantification of immunoblots of HIV-1 Cap24 protein expression normalized on Clathrin HC. (F) HIV-1 CAp24 protein quantification in supernatants assessed by ELISA and normalized to total protein amount. (G) Quantification of HIV-1 infectious particles in the supernatant using HeLa TZM bl reporter cells. (H) Quantification of lactate dehydrogenase (LDH) release in the supernatant used as a marker of cytotoxicity. (I) Relative quantity of HIV-1 total gene expression normalized on *TBP* calculated using the 2 ^(-ΔCt)^ method. Points represent the mean of 3 to 6 independent experiments +/− SEM. (C-H) One sample T-test and (I) one-way ANOVA statistical analysis were performed, the numbers above the graphs indicate the p-value when p < 0.05.

We next found that the downregulation of SNAT7 also affected the transcription of viral genes (Fig. 2I), as evidenced by the strong reduction in the expression levels of total viral genes or the gene encoding *GagPr55* (Supp. Fig. 2G).

Collectively, these results reveal that SNAT7 is crucial for HIV-1 production by human macrophages, not only at the level of viral protein synthesis, but also at the level of viral gene expression. Importantly, no global effect on transcription, translation or cell viability was detected, indicating that SNAT7 has a more specific role as a HIV-1 dependency factor.

### SNAT7 depletion alters HIV-1 reverse transcription in macrophages

To better pinpoint the stage of infection affected by SNAT7 expression, we analyzed early steps of the viral cycle. First, we quantified viral fusion at the plasma membrane using the BLaM-Vpr assay (Boncompain et al., 2019; Cavrois et al., 2002). Cells were depleted for SNAT7 or treated with control siRNA for 5 days (Fig. 3A, B).

**Figure 3.**
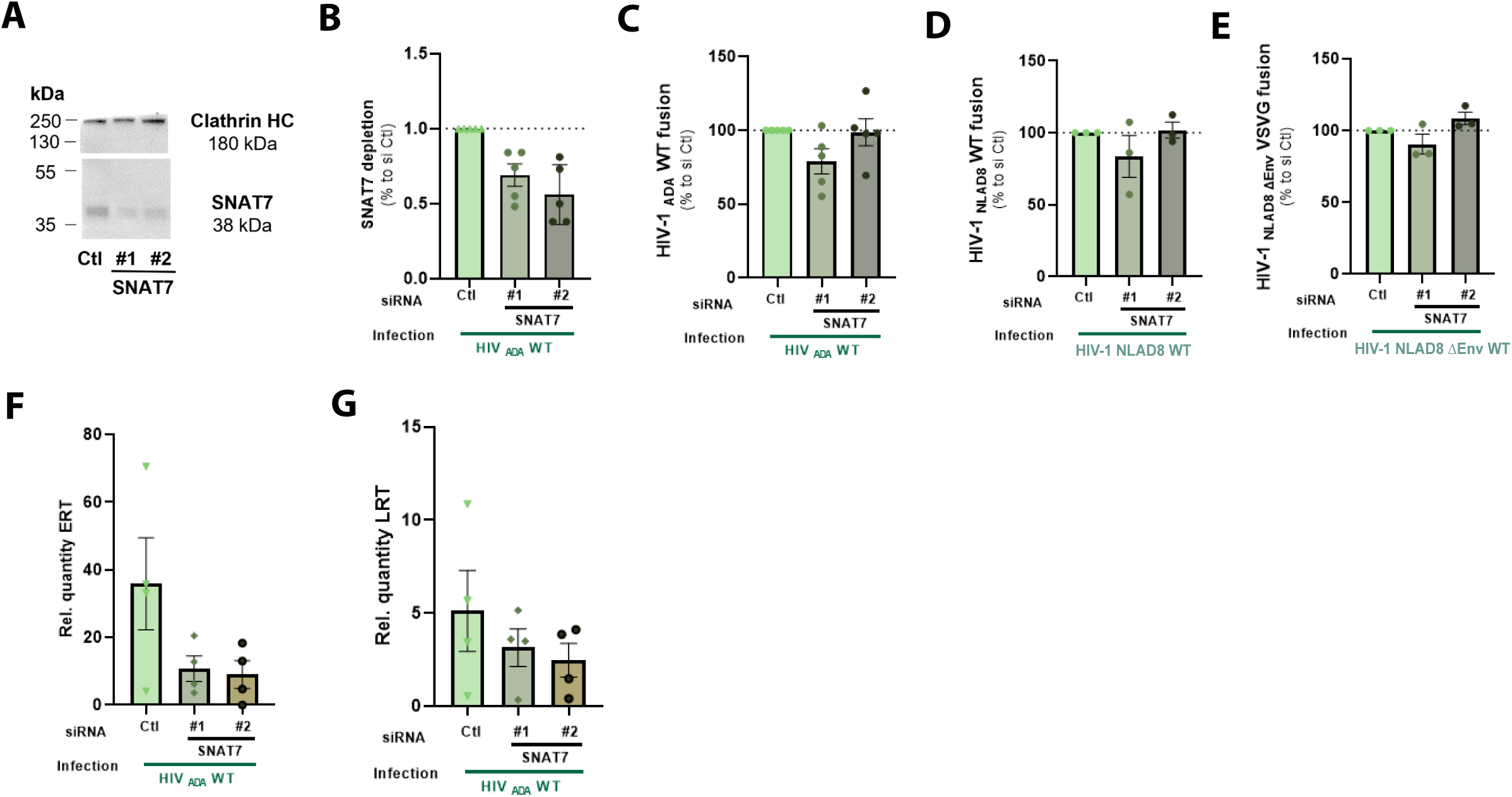
SNAT7 depletion alters reverse transcription. (A-E) Human macrophages were treated with siRNA duplexes targeting SLC38A7/SNAT7 or luciferase as control (siCtl) for 5 d. (A) Representative immunoblot of SNAT7 protein expression, in parallel with Clathrin HC as loading control, and (B) its quantification. Macrophages were infected with (C) HIV-1 ADA WT, (D) HIV-1_NLAD8_ WT and (E) VSV-G-pseudotyped HIV-1_NL4.3ΔEnv_ BLaM-Vpr-bearing particles for 1 h spinoculation at 4°C and 2.5 h infection at 37°C. Then cells were loaded with the β-lactamase substrate CCF2-AM for 2 h and fixed. Viral fusion was assessed by flow cytometry by calculating the number of cells positive for the cleaved CCF2 substrate fluorescence (447 nm) on the total amount of CCF2 positive cells (552 nm and 447 nm), and was expressed as a percentage. Points represent the mean of 5 independent experiments +/− SEM. One sample T-test statistical analysis was performed. (F-G) Human macrophages were infected during 24 h and 72 h with HIV-1 ADA WT strain, and DNA was extracted. (F) Relative quantity of HIV-1 Early Retro Transcripts (ERT) detected 24 h post-infection, normalized on *TBP*, and calculated using the 2 ^(-ΔCt)^ method. (G) Relative quantity of HIV-1 Late Retro Transcripts (LRT) detected 72 h post-infection, normalized on *TBP*, and calculated using the 2 ^(-ΔCt)^ method. Points represent the mean of 4 independent experiments +/− SEM. Ordinary one-way ANOVA statistical analysis was performed.

No signal was detected in control conditions such as uninfected cells loaded (CCF2 +) or not (CCF2 -) with CCF2-AM, or in cells infected with HIV-1_ADA_WT Vpr-Vpx-bearing particles. Pre-treatment of the cells with T-20 reduced the viral fusion events in HIV-1_ADA_ WT and _NLAD8_WT-infected cells compared to control cells, but not in VSV-G pseudotyped HIV-1_NLAD8 ΔEnv_-infected cells, as expected (Supp. Fig. 3A).

No significant decrease in viral fusion was observed in SNAT7-depleted cells compared to control cells, whether the cells were infected with HIV-1_ADA_ or _NLAD8_ strains, or VSV-G-pseudotyped particles (Fig. 3 C-E and Supp. Fig. 3A). Pre-treatment of the cells with T-20, a specific inhibitor of gp41-mediated fusion, reduced the viral fusion events in HIV-1_ADA_ WT and _NLAD8_WT-infected cells compared to control cells, but not in VSV-G pseudotyped HIV-1_NLAD8 ΔEnv_-infected cells, as expected (Supp. Fig. 3A). These results indicate that SNAT7 is not required for the viral fusion at the plasma membrane of macrophages.

We then investigated reverse transcription, a post-entry step of the viral cycle. Macrophages were depleted of SNAT7 or control siRNA for 5 days, infected with HIV-1_ADA_WT for 24 to 72 h and then lysed to collect DNA. Viral early (ERT) and late (LRT) reverse transcripts were reduced by 3.8-fold and 1.6-fold, respectively, compared to control siRNA, indicating that SNAT7 is necessary for the reverse transcription (Fig. 3F, G).

These results reveal that SNAT7 plays a role early in the viral cycle, not at the fusion-but at the reverse transcription stage.

### SNAT7 modulates the antiviral activity of SAMHD1 in macrophages

To investigate the mechanism responsible for the inhibition of the reverse transcription observed when SNAT7 is depleted, we analyzed the phosphorylation status of SAMHD1, indicative of its restriction capability, as its ability to restrict HIV-1 is negatively regulated by its phosphorylation of threonine 592 (T592) (Cribier et al., 2013; Welbourn et al., 2013; White et al., 2013). hMDM were infected or not (mock) for 8 h with HIV-1 ADA WT, washed and treated with control siRNA or SNAT7-targeting siRNA for 5 days (Fig 4A-C). As previously, the expression of GagPr55 and Cap24 decreased as a result of SNAT7 partial depletion (Supp. Fig. 4A-C). P-T592 SAMHD1 was decreased by 44 % in HIV-1-infected cells compared to control cells, in response to infection. Moreover, the partial depletion of SNAT7 by siRNA treatment induced a reduction in P-T592 SAMHD1 both in mock and HIV-1-infected cells compared to control cells (Fig. 4B and Supp. Fig. 4D), while total SAMHD1 expression was not significantly changed (Fig. 4C and Supp. Fig 4E, F).

**Figure 4.**
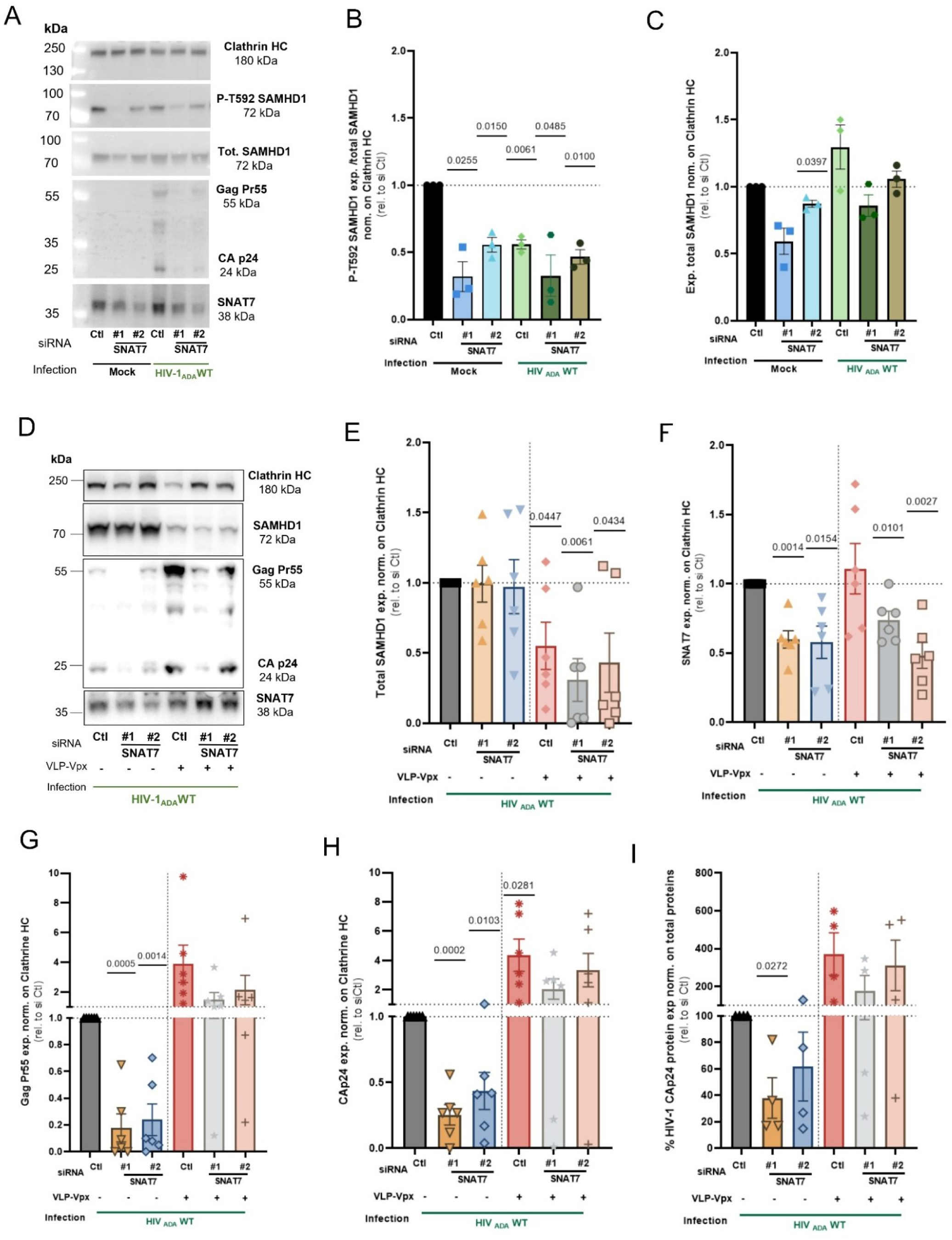
SNAT7 modulates the antiviral activity of SAMHD1. (A-C) Human macrophages were infected with the HIV-1_ADA_WT strain or mock-infected for 8 h, washed and treated with siRNA duplexes targeting SLC38A7/SNAT7 or luciferase as control (siCtl). Macrophages were cultured for 5 d, supernatants were stored and cells lysed to collect proteins. (A) Representative immunoblot of SNAT7, HIV-1 Gag Pr55 and CAp24, phospho-T592 of SAMHD1 (P-T592 SAMHD1), and total SAMHD1 protein expression, in parallel with Clathrin HC as loading control. (B) Quantification of immunoblots of P-T592 SAMHD1 protein expression normalized on the ratio of total SAMHD1 on Clathrin HC. (C) Quantification of immunoblots of total SAMHD1 protein expression normalized on Clathrin HC. The graph represents the mean of 3 independent experiments +/− SEM. One sample T-test statistical analysis was performed, the numbers above the graphs represent the p-value when p<0.05. (D-I) Human macrophages were treated with empty VLPs or VLPs bearing Vpx for 4 h, infected with the HIV-1_ADA_WT strain for 6 h, washed and treated with siRNA duplexes targeting SLC38A7/SNAT7 or luciferase as control (siCtl). Macrophages were cultured for 5 d, supernatants were stored and cells lysed to collect proteins. (D) Representative immunoblot of SNAT7, HIV-1 Gag Pr55 and CAp24, and total SAMHD1 protein expression, in parallel with Clathrin HC as loading control. (E-H) Quantification of immunoblots of (E) total SAMHD1, (F) SNAT7, (G) GagPr55 and (H) Cap24 protein expression normalized on Clathrin HC. (I) HIV-1 CAp24 protein quantification in supernatants assessed by ELISA and normalized on total protein amount. The graph represents the mean of 4 to 6 independent experiments +/− SEM. One sample T-test statistical analysis was performed.

These results suggest that SNAT7 regulates the phosphorylation status of the T592 residue of SAMHD1 and potentially its ability to restrict HIV-1 infection in human macrophages.

To further explore the link between SAMHD1 and the HIV-1 inhibition in SNAT7-depleted macrophages, we used the HIV-2 Vpx protein to degrade SAMHD1 (Ahn et al., 2012; Hofmann et al., 2012; Hrecka et al., 2011; Laguette et al., 2011). hMDM were treated with empty virus-like particles (VLP) or VLP carrying Vpx (VLP-Vpx) for 4 h, infected with HIV-1 ADA WT for 6 h, washed, and depleted for SNAT7 or with control siRNA for 5 d (Fig. 4D-I). As previously shown, SNAT7 depletion (Fig. 4D, F) induced a reduction in GagPr55 and Cap24 viral proteins production in empty VLP-treated cells (Fig. 4D, G, H, I), without affecting total SAMHD1 expression (Fig. 4D, E). As expected, Vpx delivery to cells induced SAMHD1 degradation (Fig. 4D, E) in comparison to cells treated with empty VLP. In addition, in VLP-Vpx-treated cells, GagP55 protein expression was enhanced by 3.9-fold (Fig. 4G), and Cap24 protein expression was 4.4-fold higher in siCtl cells when quantified by immunoblot (Fig. 4H) or 3.7-fold higher by ELISA (Fig. 4I), in comparison to siCtl cells treated with empty VLPs (Fig. 4G-I), confirming the restriction activity of SAMHD1 in macrophages. Interestingly, GagP55 and Cap24 proteins expression were also restored in SNAT7-depleted macrophages (Fig. 4G-I), showing that SAMHD1 is a major factor accounting for the defect in viral proteins production in the absence of SNAT7.

Taken together, these results suggest that SAMHD1 is primarily responsible for the viral restriction observed in the absence of SNAT7.

### Extracellular glutamine supplementation partially restores HIV-1 production in SNAT7-depleted macrophages

SNAT7 is the primary lysosomal transporter of glutamine (Verdon et al., 2017). To evaluate the importance of this function in HIV-1 infection of human macrophages, we assessed whether the block of infection observed in SNAT7-depleted macrophages could be reverted by adding extracellular L-glutamine to the cell cultures (Fig. 5A).

**Figure 5.**
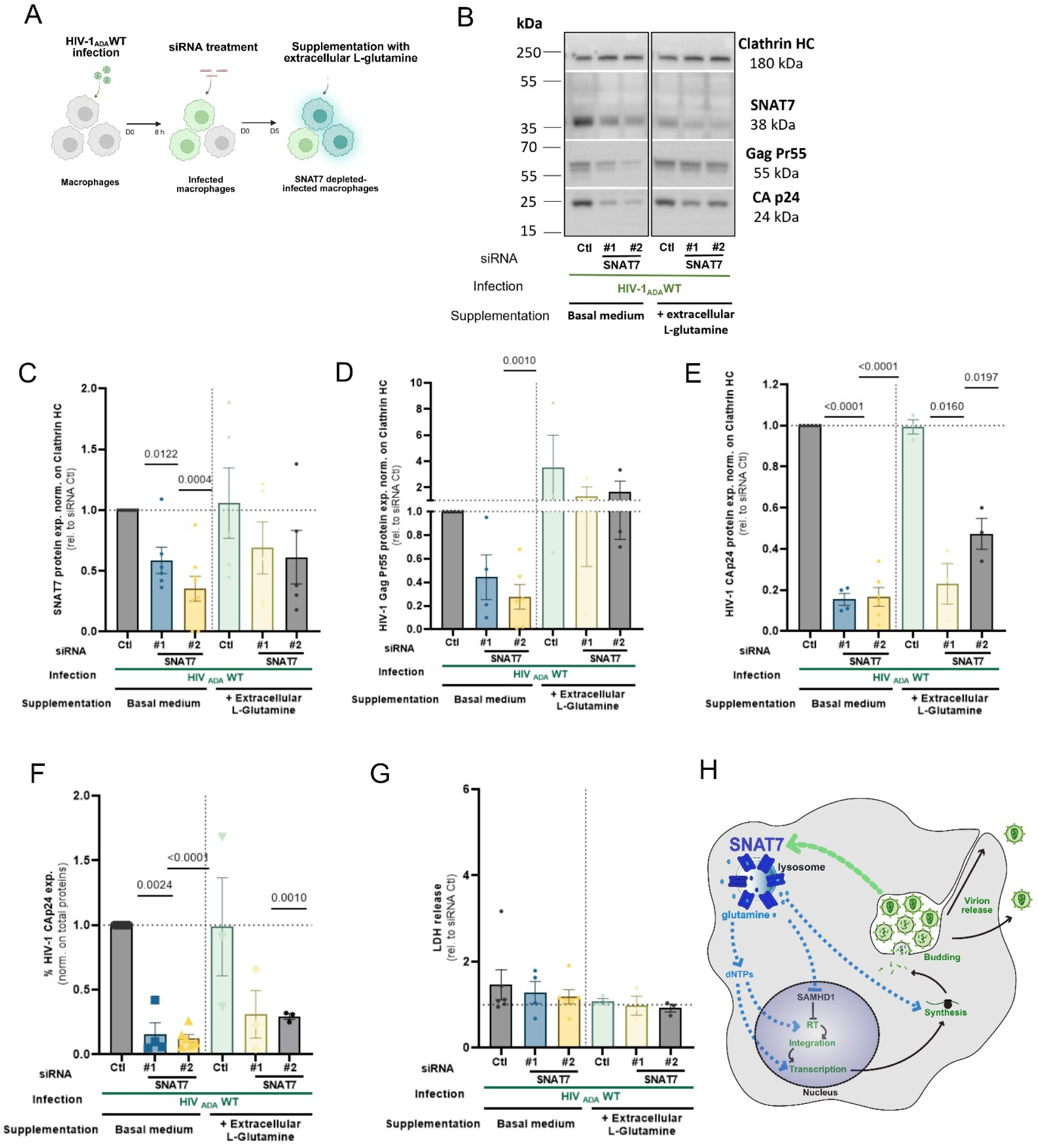
Extracellular glutamine supplementation partially restores HIV-1 production in SNAT7-depleted human macrophages. (A-G) Human macrophages were infected with the HIV-1_ADAWT_ strain or mock-infected for 8 h, washed and treated with siRNA duplexes targeting SLC38A7/SNAT7 or luciferase as control. Extracellular L-glutamine at 2 mM was added to the supernatant immediately after siRNA treatment and after 3 d of culture. Macrophages were cultured for 5 d in basal medium (i.e. without L-glutamine supplementation) or in medium supplemented with L-glutamine, supernatants were stored and cells lysed to collect proteins. (A) Experimental scheme for obtaining the biological samples analyzed in (B-G). (B) Representative immunoblot of SNAT7, HIV-1 Gag Pr55 and CAp24 protein expression, in parallel with Clathrin HC as loading control. (C) Quantification of immunoblots of SNAT7 protein expression normalized on Clathrin HC. (D) Quantification of immunoblots of HIV-1 Gag Pr55 protein expression normalized on Clathrin HC. (E) Quantification of immunoblots of HIV-1 Cap24 protein expression normalized on Clathrin HC. (F) HIV-1 CAp24 protein quantification in supernatants assessed by ELISA and normalized on total protein amount. (G) Quantification of lactate dehydrogenase (LDH) release in the supernatant used as a marker of cytotoxicity. The graphs represent the mean of 3 to 8 independent experiments +/− SEM. One sample T-test statistical analyses were performed, the numbers above the graphs represent the p-value when p < 0.05. (H) Schematic representation of how HIV-1 increases SNAT7 expression, leading to lysosomal glutamine efflux and inhibition of the anti-retroviral activity of SAMHD1, thereby promoting its own production.

Glutamine supplementation did not significantly affect SNAT7 expression (Fig. 5B, C), but partially restored HIV-1 GagPr55 expression in SNAT7-depleted cells (Fig. 5B, D-F). GagPr55 expression was enhanced by 3.5-fold in siCtl cells supplemented with glutamine, compared to siCtl cells in basal medium. Glutamine-supplemented SNAT7-depleted cells displayed 2.9-fold (for SNAT7 siRNA#1) and 5.8-fold (for SNAT7 siRNA#2) more GagPr55 protein expression than SNAT7-depleted cells cultured in basal medium, masking the impact of SNAT7 depletion (Fig. 5D). Glutamine supply however did not restore the expression of CAp24 protein to the same extent (Fig. 5E, F). Of note, none of the above-mentioned treatments altered the cell viability (Fig. 5G).

Overall, these results indicate that the supply of glutamine specifically *via* the SNAT7 transporter is involved in the production of HIV-1 GagPr55 and new virions.

Taken together, these findings highlight a novel role for glutamine delivered *via* the SNAT7 transporter in promoting viral production in human macrophages (Fig. 5H).

## Discussion

In this study, we revealed the crucial role of the lysosomal glutamine transporter SNAT7 in efficient viral infection of human macrophages. While glutamine exported to the cytosol can be directly used as a precursor for dNTP or viral proteins, the inhibition of SAMHD1 anti-retroviral activity also contributes to promote the viral production (Fig. 5H).

To characterize the role of SNAT7 in HIV-1 production, we first analyzed the later steps of the viral cycle, corresponding to the production of viral structural proteins, such as the precursor GagPr55 and the capsid p24 protein obtained from the cleavage of Gag, as well as the release of infectious particles in the supernatants of infected cells. All these events were strongly affected by the absence of SNAT7, and partially rescued by the addition of glutamine in the extracellular medium. We also observed that the transcription of viral genes was impaired when SNAT7 was downregulated. We therefore tried to determine which earlier stage of the viral cycle was affected in the absence of SNAT7. Our results showed that the viral entry in SNAT7-depleted macrophages was not modified. In contrast, the levels of viral ERT and LRT were reduced, indicating an involvement of SNAT7 in the reverse transcription stage of HIV-1. The dNTP pool is very low in hMDM and critical for HIV-1 reverse transcription (Allouch et al., 2013). Glutamine has been described as a precursor for the *de novo* biosynthesis of pyrimidines and purines (Yoo et al., 2020), which could explain the observed decrease in ERT and LRT in SNAT7-depleted macrophages.

We then focused on SAMHD1, an important restriction factor for HIV-1 that acts in particular by depleting the dNTP pool necessary for viral retro-transcription in non-cycling cells such as macrophages (Goldstone et al., 2011; Lahouassa et al., 2012) and quiescent CD4+ T cells (Baldauf et al., 2012). We assessed its antiviral activity status by monitoring its phosphorylation on the T592 residue (Cribier et al., 2013; Welbourn et al., 2013; White et al., 2013). The depletion of SNAT7 diminished SAMHD1 T592 phosphorylation. Furthermore, the degradation of SAMHD1 by Vpx induced a rescue of the viral production in SNAT7-depleted macrophages, indicating that it is a major restriction factor explaining the inhibition of viral production in the absence of SNAT7. While a direct interaction between SNAT7 and SAMHD1 cannot be excluded, we observed a partial recovery of the viral production in SNAT7-depleted macrophages cultured with extracellular glutamine supplementation, underlining the role of SNAT7 as glutamine transporter. Together, these results highlight the modulation of lysosomal glutamine efflux and the restriction factor’s activity downstream of SNAT7, but also the possibility that glutamine plays a role in the generation of nucleic acids or proteins, important pathways for efficient viral production.

The expression of the SNAT7 protein was found to be increased by the HIV-1 infection, in line with previous reports on HIV-1-induced positive or negative regulation of the expression of SLC at the protein level (Matheson et al., 2015; Montoya, 2023; Park et al., 2016). Interestingly, we noticed that several genes coding for members of the SLC family were also modulated in a transcriptomic analysis of HIV-1 infected human macrophages, together with other genes related to membrane trafficking and metabolism (Supp. Fig. 5). Of note, another member of the SLC38A family of amino acid transporters, SNAT1, an alanine transporter presents at the plasma membrane of CD4+ T cells, was shown to be modulated by Vpu in a global proteomic approach (Matheson et al., 2015). In addition, (Datta et al., 2016) reported that HIV-1 Vpr was responsible for disturbing glycolysis and glutamate metabolism in macrophages. In our experimental conditions, the absence of SNAT7 had a specific impact on the production of new HIV-1 virions. These observations might be specific to macrophages, as SNAT7 expression was low in resting monocytes and T lymphocytes, with a slight increase after T cell activation. This means that the metabolic program may be different in these cell types. Indeed, previous studies highlighted the existence of populations of CD4+ T cells with a modified metabolic status, making them more susceptible to HIV-1 infection (Clerc et al., 2019; Valle-Casuso et al., 2019).

Our results highlight SNAT7 as an important transporter needed to supply the cell with glutamine recycled from the lysosome, which is probably even more relevant for macrophages in a nutrient-restricted environment such as tissue-resident macrophage. Our findings reveal that SNAT7 is crucial for viral production, and should therefore be considered as a host-dependency factor for HIV-1, specifically in human macrophages. Given the role of these cells as viral reservoirs (Arainga et al., 2017; Castellano et al., 2017; Ganor et al., 2019; Kruize and Kootstra, 2019; Veenhuis et al., 2023; Wong et al., 2019), these results shed new light on the possibility to target membrane transporters in order to prevent viral infection.

## Acknowledgments

We wish to thank the EFS healthy donors for their generous contribution to research. We thank Philippe Benaroch (Institut Curie, Inserm U932) for early discussions on the project. We thank CYBIO and GENOM’IC facilities at Institut Cochin for their help. We thank the IMAG’IC facility of Institut Cochin that is part of the national France-BioImaging infrastructure supported by Agence Nationale de la Recherche (ANR-24-INBS-0005 FBI BIOGEN). Work in the laboratory of FN was supported by CNRS, INSERM, Université Paris Cité and grants from Fondation pour la Recherche Médicale (FRM DEQ20130326518) and ANRS (AAP2015-2). All schemes were created with BioRender.com and Affinity Designer.

## Author contributions

Conceptualization: FH, GLB and FN; Data curation: FH, GLB; Formal analysis: FH, GLB; Funding acquisition: FN; Investigation: FH, GLB, QP, DTP, JH, TMS and AD; Methodology: FH, GLB, AD, and FN; Project administration: FH and FN; Resources: MM and FMG; Supervision: CS, ASC and FN; Validation: FH, GLB, and FN; Visualization: FH and FN; Writing – original draft: FH and FN; Writing – review & editing: FH, FN, FMG, CS and ASC.

## Methods

### Cell culture

Fresh blood was collected from healthy donors from Etablissement Français du Sang Ile-de-France, Site Trinité (Inserm agreement #15/EFS/012, #18/EFS/030 and CNRS No.2023-2026-025 CCPSL CNRS, ensuring that all donors gave a written informed consent and providing anonymized samples). The procedures for sample collection and conservation were declared and approved through CODECOH (COnservation d’Eléments du COrps Humain, No. DC-2021-4166) by the French Ministry of Higher Education, Research and Innovation, in accordance with French regulations on human biological elements. Human monocytes were isolated by 2 successive sedimentation steps using density gradients in Ficoll®-Paque Plus and then Percoll® (Merck/Cytiva™).

To obtain a purer population of T lymphocytes and monocytes for certain procedures, cells were negatively isolated from peripheral blood mononuclear cells (PBMCs) after the Ficoll separation step using EasySep™ human T cell and EasySep™ human monocytes enrichment kits (STEMCELL™ Technologies).

Isolated monocytes were cultured and differentiated in human monocyte-derived macrophages (hMDM) for 6 to 10 days in complete culture medium (RPMI 1640 supplemented with FCS 10 %, streptomycin/penicillin 100 µg/ml, and L-glutamine 2 mM (Life Technologies™/Gibco) or R10 medium (RPMI 1640 supplemented with FCS 10 %, HEPES 10 mM, sodium pyruvate 1 mM, non-essential amino acids 1 X and streptomycin/penicillin 100 µg/ml (Life Technologies™/Gibco)) containing 10 ng/ml of recombinant human (rh) M-CSF (R&D Systems), or 0.5 ng/ml of rhM-CSF and 4 ng/ml of rhGM-CSF (Miltenyi Biotech). The culture medium was then changed using complete culture medium.

Lymphocytes were activated with IL2 300 U/ml (Miltenyi Biotech) and PHA-P 1 µg/ml (Merck) from day 0 to day 3 post-isolation, and then only with IL2 for 3 more days.

### siRNA treatment

Small interfering RNA (siRNA) duplexes were diluted in OptiMEM, and mixed with Lipofectamine™ RNAiMax reagent according to the manufacturer’s instructions (Life Technologies™/Gibco) at room temperature (RT) for 20 min. The complexes were added to hMDM at a final concentration of 240 nM, and the cells were cultured for 5 to 6 d at 37°C.

The siRNA sequences were as follows: SLC38A7/SNAT7 sequence #1: 5’-CAGGUCUAAUGUUUACAAACGGUGC-3’, SLC38A7/SNAT7 sequence #2: 5’- CCAUAGCUAAUAGACAUUUCCCAGG-3’, SLC1A2 sequence #1: 5’-AUGAACGUCUUAGGUCUGAUA-3’, SLC1A2 sequence #2: 5’-CACGAGAGCCAUGGUGUAUUA-3’. The control siRNA sequence targeting luciferase was: 5’-CGUACGCGGAAUACUUCGA-3’.

### Viral production and hMDM infection

HIV-1 virions with a natural envelope or pseudotyped with VSV-G, and VSV-G pseudo-particles containing BLaM-Vpr or Vpr-Vpx were obtained by transient co-transfection of HEK293T cell line (Human Embryonic Kidney 293, ATCC®CRL1573™) using FuGENE® 6 Transfection Reagent (Promega) as recommended by the manufacturer, with the corresponding plasmids: pHIV-1_ADA_WT was kindly provided by Luciana da Costa ((Federal University); (Dumas et al., 2015; Mazzolini et al., 2010)), pHIV-1_NLAD8_WT was kindly provided by S. Benichou (Institut Cochin, Paris), pHIV-1_NL4.3_ΔEnv was a kind gift from G. Pancino (Institut Pasteur, Paris), pCMV-BlaM-Vpr encoding β-lactamase fused to the viral protein Vpr was described in (Cavrois et al., 2002), pMD.G encoding VSV-G, pVpr-Vpx, and pAdvantage.

Supernatants from transfected cells were collected after 48 h, and filtered on a 0.45 µm membrane. Supernatants containing pseudotyped particles carrying BLaM-Vpr or Vpr-Vpx were ultracentrifuged at 60 000 g for 90 min at 4°C on a sucrose cushion. All virus stocks were stored at -80°C.

SIV Viral-Like Particles (VLPs) “Cis” (Vpx expressed from the packaging vector) were produced in 293T by the calcium-phosphate co-precipitation method. VLPs were produced by transfection of VSV-G expression vector and SIV3+ΔVpr packaging vector (VLPs Vpx) or SIV3+ΔVprΔVpx (VLP control). All packaging vector were a gift from N. Landau and is described in (Gramberg et al., 2010). Cell culture medium was collected 48 h after transfection and filtered through 0.45 μm pore filters. Viral particles were concentrated by sucrose gradient and ultracentrifugation. The incorporation of Vpx into VLPs was assessed by Western-Blot as described here (Chougui et al., 2018).

Quantification of HIV-1 infectious particles was determined using HeLa-CD4 TZM-bl cells bearing the β-galactosidase gene under the control of HIV-1 LTR (NIH reagent program) as described in (Clavel and Charneau, 1994), by scoring β-lactamase-positive cells 24 h after infection.

The amount of HIV-1 p24 antigen was quantified using an HIV-1 p24 ELISA kit (Revvity/PerkinElmer) according to the manufacturer’s recommendations. The amount of p24 protein was normalised to the amount of total cellular protein.

hMDM were pre-treated or not with 100 ng of control VLPs or Vpx-containing VLPs for 4 h at 37°C and/or infected with HIV-1 strains at a multiplicity of infection (MOI) of 0.5 or treated with supernatant of HEK293T without HIV-1 particles (Mock), and incubated at 37°C for 8 h or 48 h. Cells were washed, treated with siRNA and/or cultured in fresh medium during the specified time of each experiment. The macrophage culture medium was supplemented occasionally with L-glutamine 2 mM (Gibco) for several days.

For the BLaM-Vpr fusion assay, hMDMs were pre-treated or not for 30 min on ice with 5 µM of T-20 fusion inhibitor (Kilby J. M. et al., 1998) as negative control for the test, kindly provided by J. Dutrieux, Institut Cochin, Paris,. Macrophages were then brought into contact with the BLaM-Vpr or Vpr-Vpx containing particles, spinoculated during 1 h at 1 770 g, 4°C, and incubated for 2.5 h at 37°C. Macrophages were loaded or not with the CCF2-AM substrate (Life Technologies™) of the β-lactamase for 2 h, and fixed using PFA 4 %. Enzymatic cleavage of CCF2/AM (Cavrois et al., 2002) was assessed by flow cytometry (LSR II, BD). The fusion score was determined by calculating the ratio of the number of cells positive for the cleaved CCF2-AM substrate fluorescence (447 nm) to the total number of CCF2-AM positive cells (552 nm and 447 nm) (Boncompain et al., 2019).

### Immuno-blot analysis

Cells were lysed on ice during 15 min using an ice-cold buffer containing Tris pH 7.5 20 mM, NaCl 150 mM, NP40 0.5 %, sodium fluoride (NaF) 50 mM, sodium orthovanadate 1 mM and a cOmplete™ Protease Inhibitors cocktail tablet (Merck/Roche). Samples were centrifuged at 10 000 g, 4°C for 10 min. Post-nuclear supernatants were stored on ice and quantified using the Pierce™ BCA protein assay (Life Technologies™). The same amount of protein for each condition was aliquoted and denatured at 70°C for 10 min in protein loading buffer containing Tris pH 6,8 0.31 M, glycerol 50 %, DTT 100 mM, SDS 10 %, and bromophenol blue. Cell lysates were separated by electrophoresis on 4-12 % gradient NuPAGE Bolt Bis-Tris Plus gel (Life Technologies™), and transferred to PVDF membrane (Merck/Millipore).

Membranes were incubated with diluted primary antibodies in 5 % non-fat milk/PBS 1 X, or 5 % BSA/TBS 1 X, 0.1 % Tween® 20 at 4°C with gentle shaking, overnight: rabbit anti-human SLC38A7 (Merck/Atlas Antibodies), mouse anti-clathrin heavy chain (BD Biosciences), mouse anti-vinculin (Merck), mouse anti-SAMHD1 (clone OTI1F9, Abcam), rabbit anti-phospho-SAMHD1 (Thr592, clone D7O2M, Ozyme) (Dragin L., 2013), mouse anti-HIV-1 GagPr55/Cap24 (gift from P. Benaroch, Institut Curie, Paris). After washing, membranes were incubated with secondary antibodies anti-mouse and/or anti-rabbit IgG coupled to HRP (Jackson ImmunoResearch). Protein detection was performed using SuperSignal™ West Dura ECL substrate (Life Technologies™). Chemiluminescence was captured by a 16-bit camera (Fusion FX, Vilber), and quantified using ImageJ software.

### RT-qPCR

Cells were washed twice with PBS 1 X, lysed, and RNA or DNA were extracted using the RNeasy extraction kit or the DNeasy Blood & Tissue Kit respectively (Qiagen) according to the manufacturer’s instructions. RNA concentration was quantified using the NanoDrop (Mettler Toledo), and 1 µg of mRNA was reverse transcribed in DNA using the SuperScript II Reverse Transcriptase (ThermoFisher). qPCR was performed using the LightCycler 480 SYBR Green I Master (Merck/Roche), with specific oligos to detect: TBP (Forw.: GAGCCAAGAGTGAAGAACAGTC, Rev.: GCTCCCCACCATATTCTGAATCT) used as loading control, and SNAT7 (Forw.: TCATAGCCCTCGCTGTCTACA, Rev.: GAGCACGCTCAGGATGATGAA) were designed on Primer Bank website; HIV-1 Gag (Forw.: GTGTGGAAAATCTCTAGCAGTGG ; Rev.: CGCTCTCGCACCCATCTC) from (Wang et al., 2019) ; HIV-1 total mRNA (Forw.: TTGCTTAATGCCACAGCTAT, Rev.: TTTGACCACTTGCCACCCAT) were designed with the kind help of S. Gallois-Montbrun, Institut Cochin, Paris; ERT (Forw. “NEC152”: GCCTCAATAAAGCTTGCCTTGA, Rev. “NEC131”: GGCGCCACTGCTAGAGATTTT); LRT (Forw. “F592”: CTTTCGCTTTCAAGTCCCTGTT, Rev. “R666”: AGATCCCTCAGACCCTTTTAGTCA), and albumin (Forw.: TGCATGAGAAAACGCCAGTAA, Rev.: ATGGTCGCCTGTTCACCAA) were adapted from (Allouch et al., 2013). The fold change in gene expression was assessed by calculating the 2 ^(-ΔΔCt)^ method, while the relative quantity was determined by calculating the 2 ^(-ΔCt)^ method.

### Transcriptomic analysis

hMDMs from 3 different healthy donors were differentiated for 11 d as described above and then infected or not with the HIV-1 R5-tropic strain ADA WT (HIV-_1ADA_ WT) for 7 d. RNA was extracted using RNeasy® Plus mini kit (Qiagen) according to the manufacturer’s instructions. After quality control (Bioanalyseur 2100 Agilent RNA6000 nano chip kit), 50 ng of total RNA were reverse transcribed following the Ovation PicoSL WTA System V2 (Nugen). The resulting double strand cDNA was used for amplification based on SPIA technology.

After purification using the Nugen protocol, 2.5 µg of single-stranded DNA was used for fragmentation and biotin labelling using Encore Biotin Module (Nugen). After fragmentation control using Bioanalyzer 2100, cDNA was hybridized to GeneChip® human Gene 1.0 ST (Affymetrix) at 45°C for 17 h. Chips were then washed on the fluidic station FS450 according to specific protocols (Affymetrix) and scanned using the GCS3000 7G. The image was then analyzed using the Expression Console software (Affymetrix) to obtain raw data (cel files) and metrics for Quality Controls. The observations of some of these metrics and the study of the distribution of raw data showed no outlier experiment. Data were normalized using the RMA algorithm in Bioconductor with the custom CDF vs 16 (Nucleic Acid Research 33 (20), e175). Statistical analysis was carried out with the use of Partek® GS. All data have been deposited in NCBI’s Gene Expression Omnibus and are accessible through GEO Series accession number GSE272009 (https://www.ncbi.nlm.nih.gov/geo/query/acc.cgi?acc=GSE272009).

First, variations in gene expression were analyzed using unsupervised hierarchical clustering and PCA to assess data from technical bias and outlier samples. To find differentially expressed genes (DEG), a three-ways ANOVA analysis was applied for each gene, and made pair wise Tukey’s post hoc tests between groups. Then, p-values and fold changes to filter and select DEG were used. Heatmaps were generated using R software with “pheatmap” function on normalized data, then clusters were extracted with “rect.hclust” function. DEGs enrichment analyses were carried out through the use of DAVID Bioinformatics Resources (Huang et al., 2009) for Gene Ontology analysis (Biological Process) of extracted clusters from heatmap.

### Cytotoxicity

Cellular cytotoxicity was determined by quantification of lactate dehydrogenase (LDH) in the hMDM supernatant, using the CyQUANT™ LDH kit (Life Technologies™) according to the manufacturer’s instructions.

## Statistical analysis

Statistical significance of the data was evaluated by appropriate tests (one-way or two-way ANOVA and one-sample t-test) using GraphPad Prism 9.5 software. Differences were considered significant when the p-value was < 0.05.

---

**Supplementary Figure 1.**
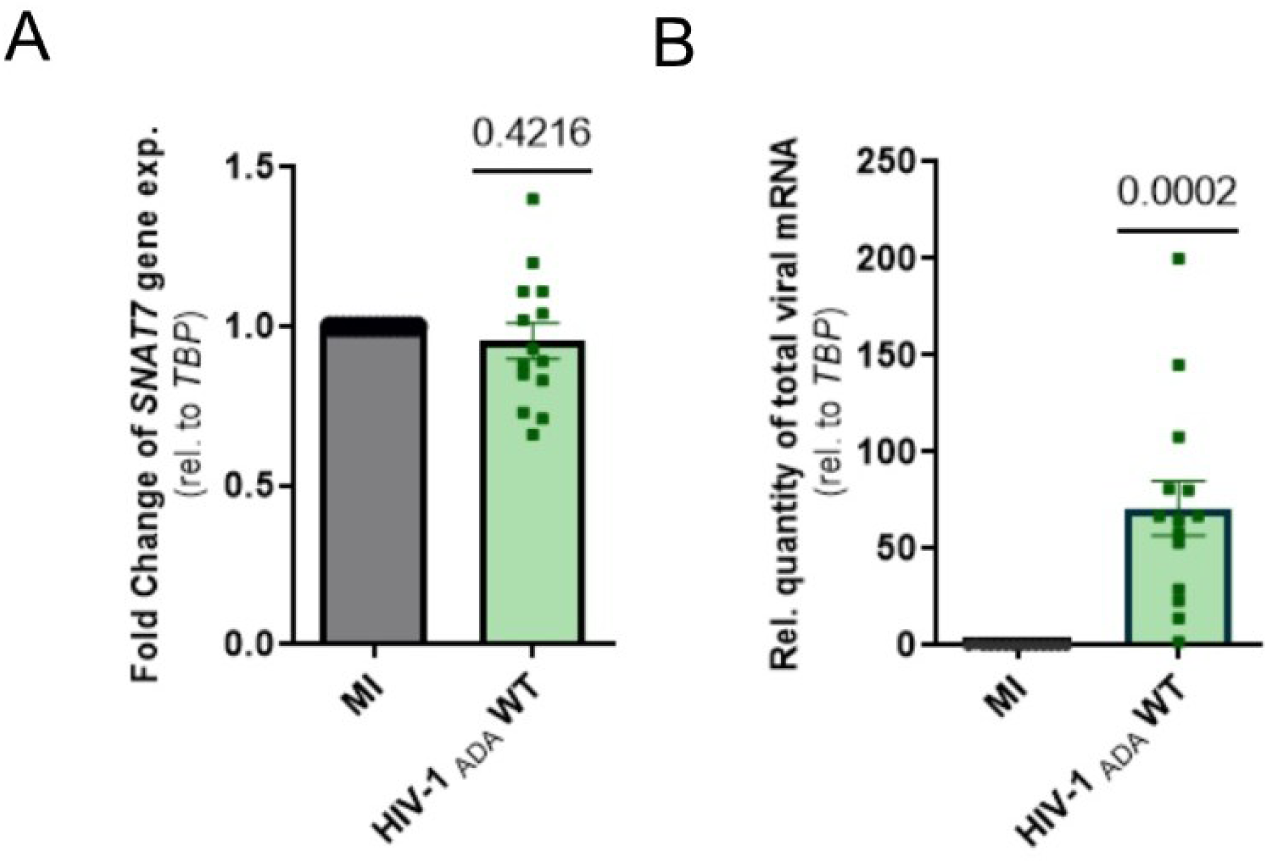
The lysosomal transporter of glutamine SLC38A7/SNAT7 is predominantly expressed by human macrophages, and its protein expression is increased upon HIV-1 infection. (A, B) Macrophages were infected with the HIV-1_ADA_WT strain or mock-infected for 6 d, and cells were lysed to collect RNA. (A) Fold change in *SNAT7* gene expression normalized on *TBP* calculated using the 2 ^(-ΔΔCt)^ method. (B) Relative quantity of HIV-1 gene expression normalized on *TBP* calculated using the 2 ^(-ΔCt)^ method. Points represent the mean of 14 independent experiments +/− SEM. One sample T-test statistical analysis was performed, the numbers above the graphs represent the p-value. (C-F) Primary human macrophages were infected with HIV-1 strain ADA WT for 6 d, with a MOI of 0.05, 0.1, 0.2 or 0.5, or were treated with supernatant from uninfected cells (Mock). The cells were then lysed and the proteins collected. (C) Representative immunoblot of GagPr55, CAp24 and SNAT7, in parallel with Clathrin Heavy Chain (Clathrin HC) used as a loading control. (D) Quantification of SNAT7 protein expression normalized to Clathrin HC. Results are expressed as a ratio of protein expression to the control condition (Mock). (E, F) Quantification of GagPr55 and CAp24 protein expression normalized to Clathrin HC. Results are expressed as a ratio of protein expression to MOI 0.05 condition. Graphs represent the mean of 3 independent experiments +/- SEM. A non-parametric Kruskal-Wallis ANOVA statistical test was performed.

**Supplementary Figure 2.**
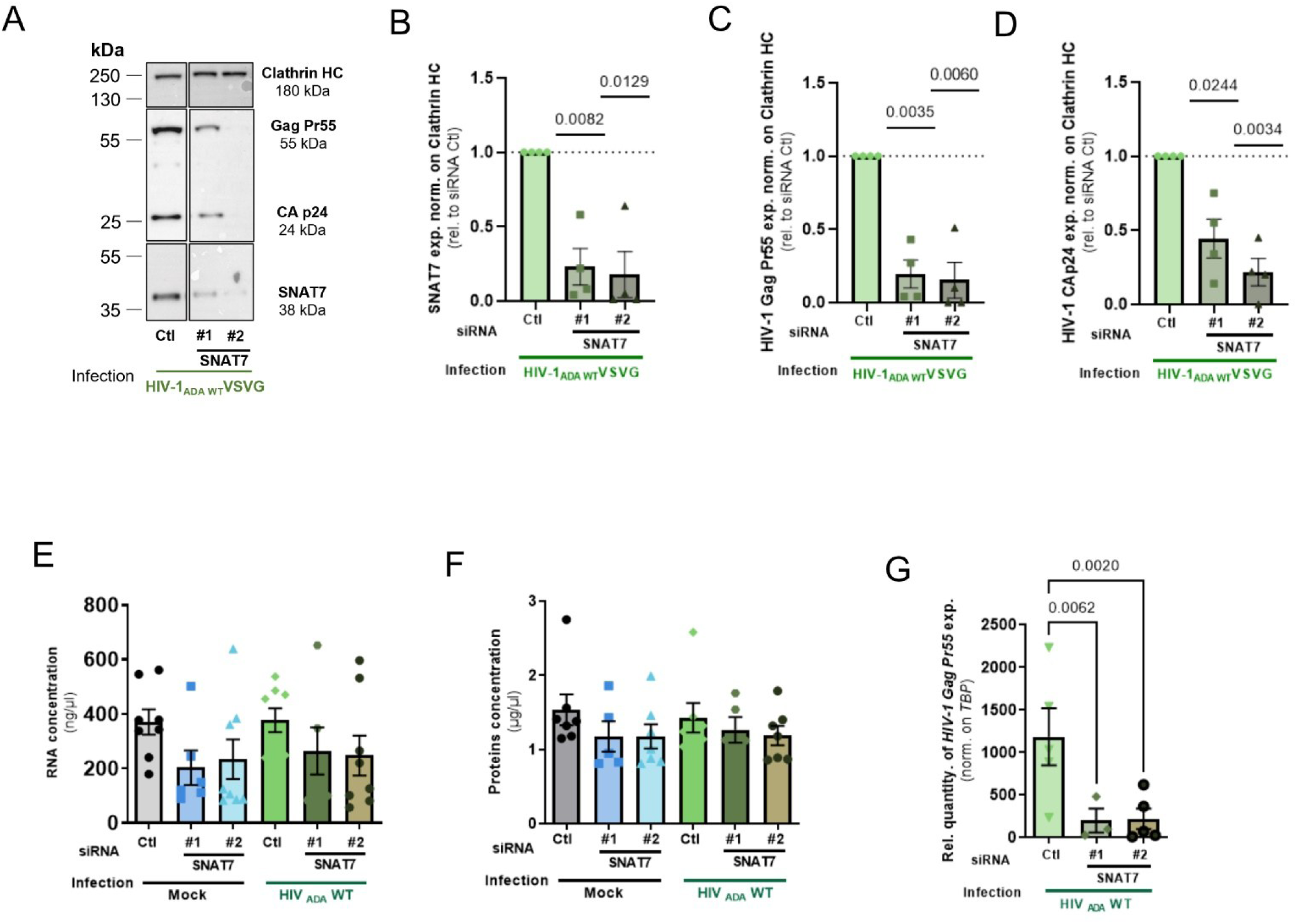
SNAT7 is necessary for the efficient production of HIV-1 by human macrophages. (A-D) Human macrophages were infected with the HIV-1ADA WTVSVG strain or mock-infected for 8 h, washed and treated with siRNA duplexes targeting SLC38A7/SNAT7 or luciferase as control. Macrophages were cultured for 5 d, and cells lysed to collect proteins. (A) Representative immunoblot of SNAT7, HIV-1 Gag Pr55 and CAp24 protein expression, in parallel with Clathrin HC as a loading control. (B) Quantification of immunoblots of SNAT7 protein expression normalized to Clathrin HC. (C) Quantification of immunoblots of HIV-1 Gag Pr55 protein expression normalized to Clathrin HC. (D) Quantification of immunoblots of HIV-1 Cap24 protein expression normalized on Clathrin HC. Points represent the mean of 4 independent experiments +/− SEM. One sample T-test statistical analysis was performed, the numbers above the graphs indicate the p-value when p < 0.05. (E-G) Human macrophages were infected with the HIV-1_ADA_WT strain or mock-infected for 8 h, washed and treated with siRNA duplexes targeting SLC38A7/SNAT7 or luciferase as control. Macrophages were cultured for 5 d, and cells lysed to collect RNA and proteins. Concentration of total (A) RNA and (B) proteins extracted from cells. (C) Relative quantity of HIV-1 GagPr55 gene expression normalized on TBP calculated using the 2 (-ΔCt) method. Points represent the mean of 3 to 8 independent experiments +/− SEM. One-way ANOVA statistical analysis was performed, the numbers above the graphs represent the p-value when p < 0.05.

**Supplementary Figure 3.**
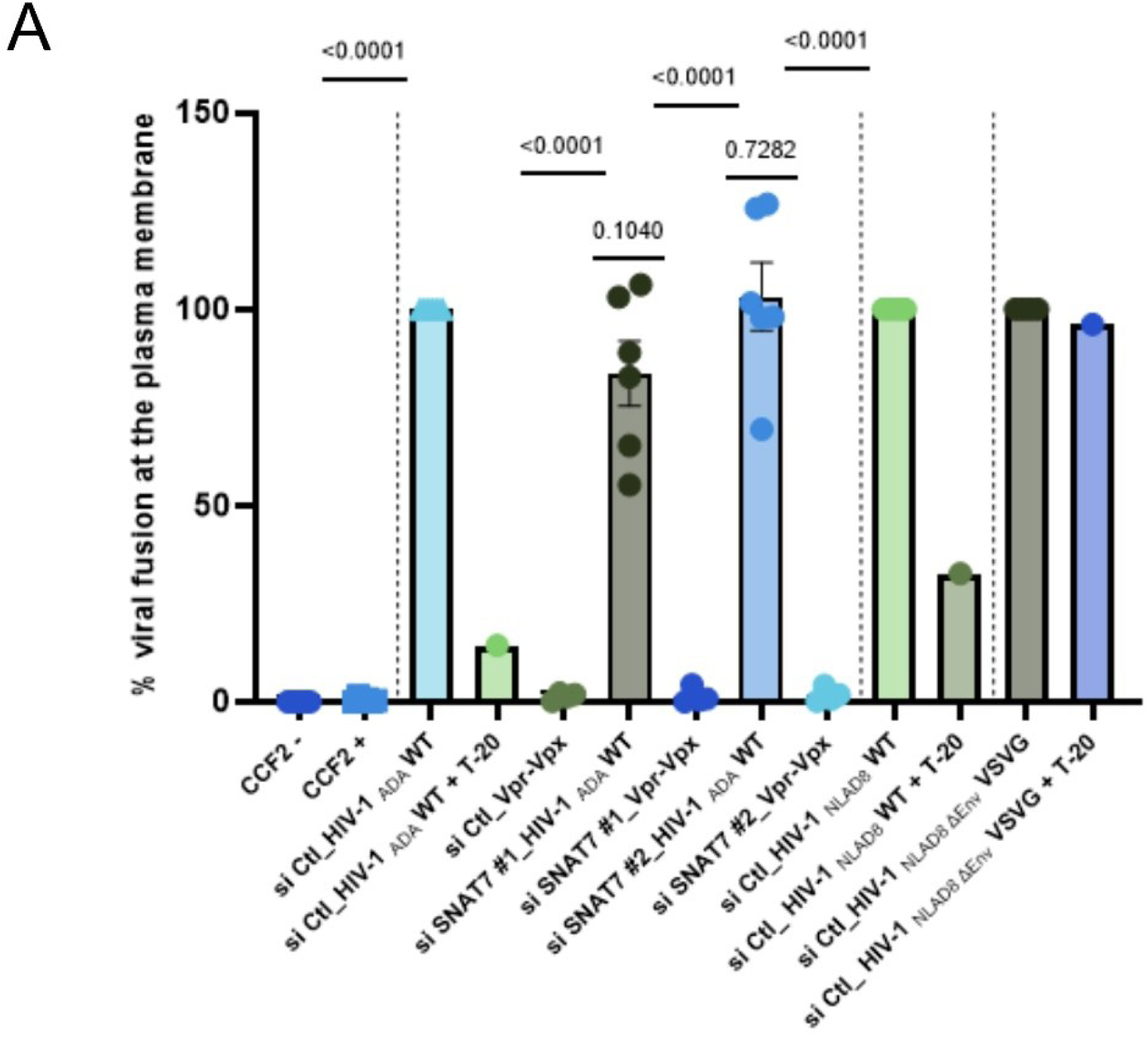
SNAT7 depletion alters reverse transcription. (A) Human macrophages were treated with siRNA duplexes targeting SLC38A7/SNAT7 or luciferase as control. Macrophages were cultured for 5 d, pre-treated or not with the fusion inhibitor T-20 at 5 µM for 30 min on ice, and infected or not (CCF2 – and +) with HIV-1_ADA_WT, HIV-1_NLAD8_WT and VSV-G-pseudotyped HIV-1_NL4.3ΔEnv_ BLaM-Vpr-bearing particles or HIV-1_ADA_WT Vpr-Vpx-bearing particles for 1 h spinoculation at 4°C, and 2.5 h culture at 37°C. Then cells were loaded or not (CCF2 -) with the β-lactamase substrate CCF2-AM for 2 h and fixed. Viral fusion at the plasma membrane was assessed by flow cytometry by calculating the number of cells positive for the cleaved CCF2-AM substrate fluorescence (447 nm) on the total amount of CCF2-AM positive cells (552 nm and 447 nm), and was expressed as a percentage. Points represent the mean of 4 to 5 independent experiments +/− SEM, except for T-20 treatment (n = 1). One sample T-test statistical analysis was performed, the numbers above the graphs represent the p-value when p<0.05.

**Supplementary Figure 4.**
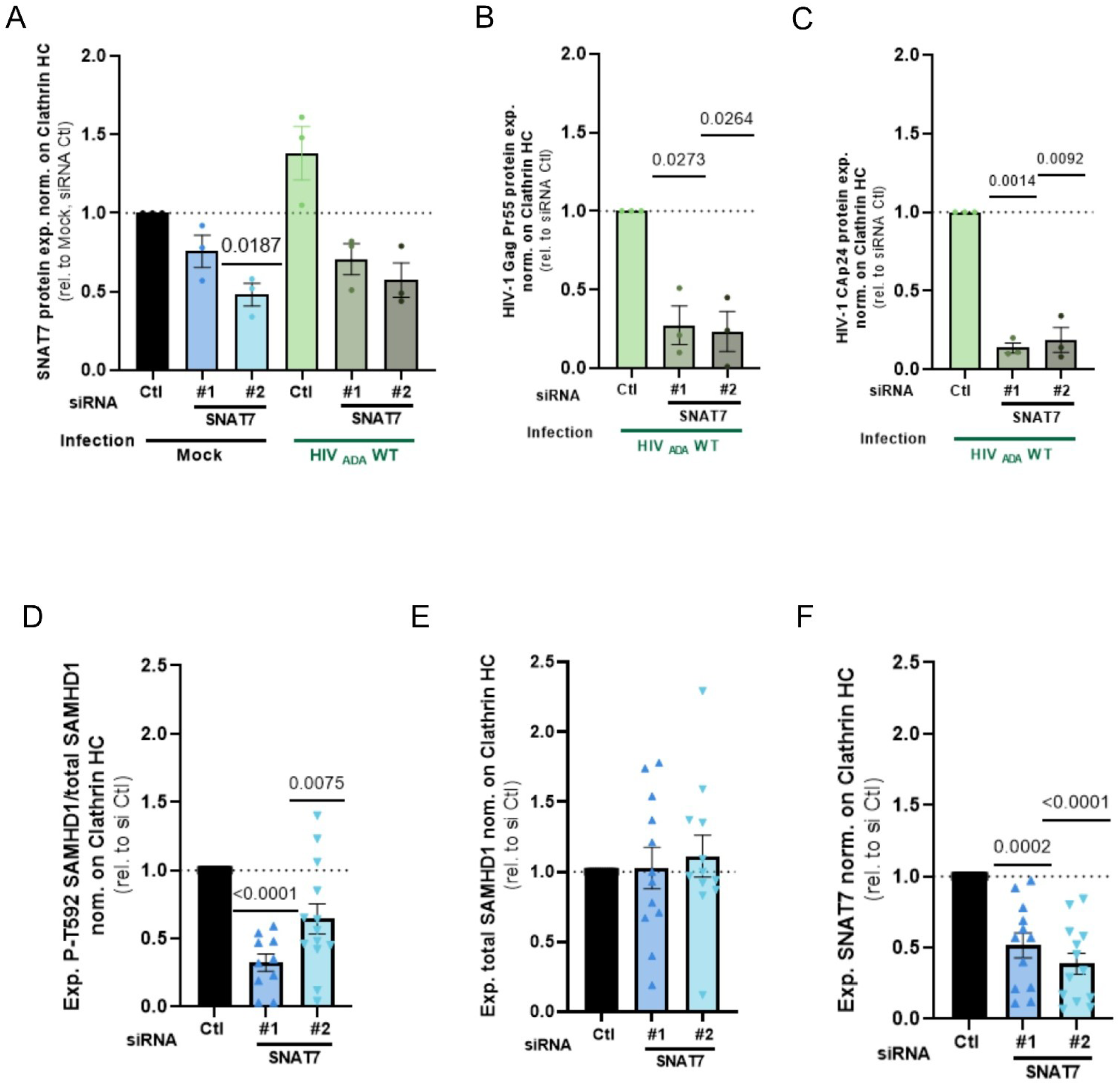
SNAT7 modulates the antiviral activity of SAMHD1. (A-C) Human macrophages were infected with the HIV-1_ADA_WT strain or mock-infected for 8 h, washed and treated with siRNA duplexes targeting SLC38A7/SNAT7 or luciferase as control (siCtl). Macrophages were cultured for 5 d and cells lysed to collect proteins. Quantification of immunoblots of (A) SNAT7, (B) GagPr55 and (C) Cap24 protein expression normalized on Clathrin HC. The graph represents the mean of 3 independent experiments +/− SEM. One sample T-test statistical analysis was performed, the numbers above the graphs represent the p-value when p<0.05. (D-F) Human macrophages were treated with siRNA duplexes targeting SLC38A7/SNAT7 or luciferase as control. Macrophages were cultured for 5 d, and cells lysed to collect proteins. (D) Quantification of immunoblots of P-T592 SAMHD1 protein expression normalized on the ratio of expression of total SAMHD1 on Clathrin HC. (E) Quantification of immunoblots of total SAMHD1 protein expression normalized on Clathrin HC. (F) Quantification of immunoblots of SNAT7 protein expression normalized on Clathrin HC. Points represent the mean of 13 independent experiments +/− SEM. One sample T-test statistical analysis was performed, the numbers above the graphs represent the p-value when p<0.05.

**Supplementary Figure 5.**
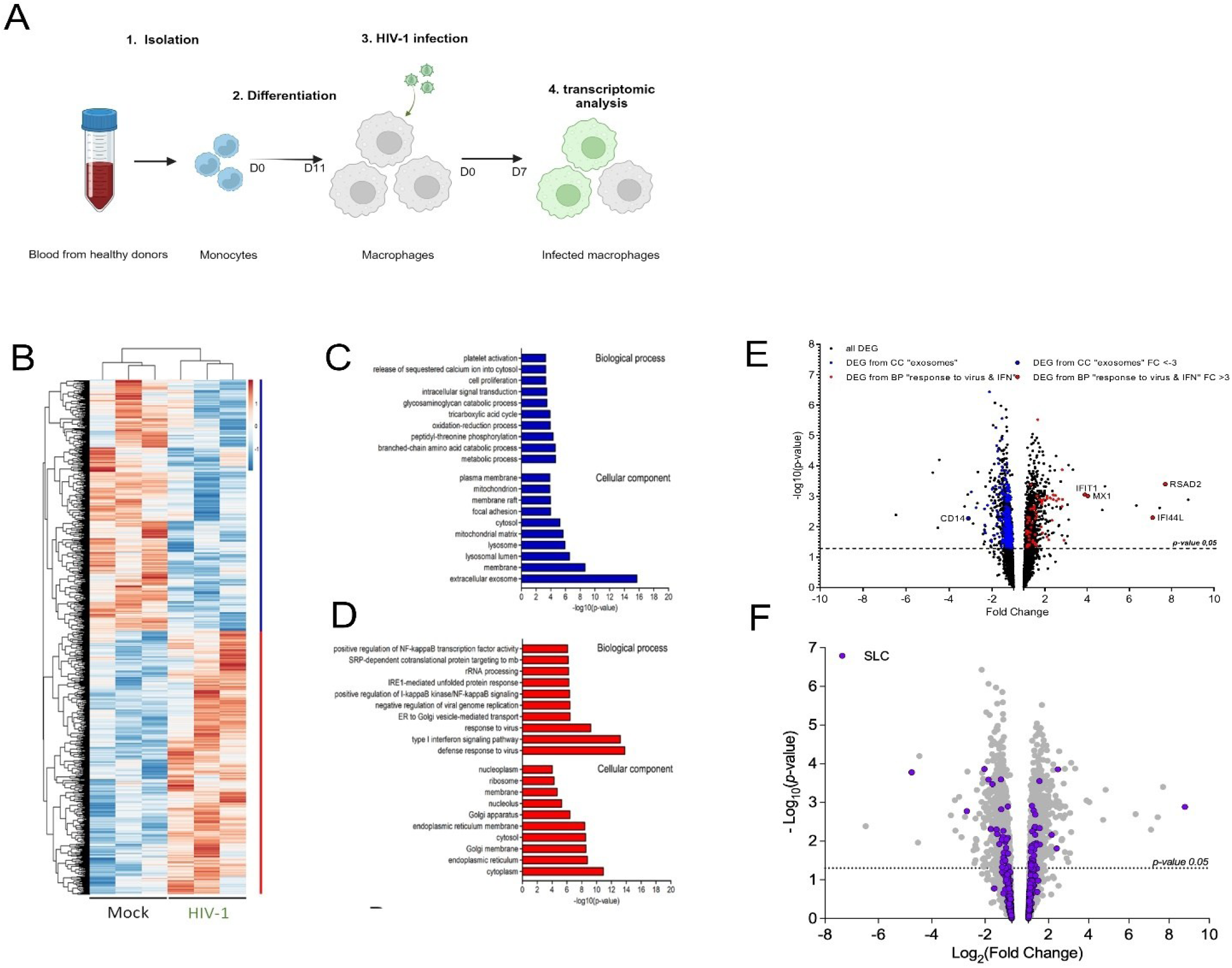
The expression of SLC genes is modulated by HIV-1 infection in human macrophages. (A-F) Human monocytes were isolated from the blood of 3 different donors and differentiated into macrophages for 11 d. Cells were infected with the HIV-1 macrophage-tropic strain ADA WT for 7 d and lysed to extract RNA and perform a transcriptomic analysis. (A) Experimental scheme showing how the biological samples analyzed were obtained in (B-F). After reverse transcription, cDNA was analyzed on GeneChip® human Gene 1.0 ST (Affymetrix). After statistical analysis, heat maps were built (B) and negatively (C) or positively (D) modulated gene clusters were extracted using R software. Differentially expressed genes (DEGs) enrichment analysis have been achieved through the use of DAVID Bioinformatics Resources for Gene Ontology analysis (Biological Process/Cellular Component) of extracted clusters from the heat map (E-F)

